# DeepMIB: User-friendly and open-source software for training of deep learning network for biological image segmentation

**DOI:** 10.1101/2020.07.13.200105

**Authors:** Ilya Belevich, Eija Jokitalo

**Affiliations:** Electron Microscopy Unit, Institute of Biotechnology, University of Helsinki, PO Box 56, FI-00014 Helsinki, Finland

## Abstract

Deep learning approaches are highly sought after solutions for coping with large amounts of collected datasets and are expected to become an essential part of imaging workflows. However, in most cases, deep learning is still considered as a complex task that only image analysis experts can master. DeepMIB addresses this problem and provides the community with a user-friendly and open-source tool to train convolutional neural networks and apply them to segment 2D and 3D light and electron microscopy datasets.

## Introduction

During recent years, improved availability of high-performance computing resources, and especially graphics processing units (GPUs), has boosted applications of deep learning techniques into many aspects of our lives. The biological imaging is not an exception, and methods based on deep learning techniques [1] are continually emerging to deal with various tasks such as image classification [2, 3], restoration [4], segmentation [5–8], and tracking [9]. Unfortunately, in many cases, the application of these methods is not easy and requires significant knowledge in computer sciences, making it difficult to adapt by many researchers. Software developers have already started to address this challenge by developing user-friendly deep learning tools, such as Cell Profiler [10], Ilastik [11], ImageJ plug-ins DeepImageJ [12] and U-net [13], CDeep3M [5], and Uni-EM [14] that are especially suitable for biological projects. However, the overall usability is limited because they either rely on pre-trained networks without the possibility of training on new data [10–12], are limited to electron microscopy (EM) datasets [14], or have specialized computing requirements [5, 13].

In our opinion, in addition to providing good results, ideal deep learning solution should fulfill the following criteria: a) is capable of training on new data; b) has a user-friendly interface; c) is easy to install and would work straight out of the box; d) is free of charge and e) has open-source code for future development. Preferably, it also would be compatible with 2D an 3D, EM and light microscopy (LM) datasets. To address all these points, we are presenting DeepMIB as a free open-source software tool for image segmentation using deep learning. DeepMIB can be used to train 2D and 3D convolutional neural networks (CNN) on user’s isotropic and anisotropic EM or LM multicolor datasets. DeepMIB comes bundled with Microscopy Image Browser (MIB) [15], forming a powerful combination to address all aspects of an imaging pipeline starting from basic processing of images (*e.g.*, filtering, normalization, alignment) to manual, semi-automatic and fully automatic segmentation, proofreading of segmentations, their quantitation and visualization.

## Software availability

The software is distributed either as an open-source Matlab code or as a compiled standalone application for Windows and MacOS. DeepMIB is automatically installed during installation of MIB (version 2.70 or newer). The Matlab version requires license for Matlab and Deep Learning, Computer Vision, Image Processing toolboxes. The compiled versions do not require Matlab license and are installed using the provided installer. All distributions and installation instructions are available directly from MIB website (http://mib.helsinki.fi) or from GitHub (https://github.com/Ajaxels/MIB2). All presented examples below can be downloaded and tested (see Supplementary Material for details).

## Application of DeepMIB to LM and EM datasets

We have tested DeepMIB on several 2D and 3D datasets from both LM and EM, and present here examples from each of the four types (Fig. 1, Supplementary Movie S1-4). These results should not be considered as winners of a segmentation challenge but rather as examples of what inexperienced users can achieve on their own with only basic knowledge of CNNs. The first example demonstrates the segmentation of membranes from a 2D-EM image, which can be individual micrographs of thin TEM sections or slices of 3D-datasets. Here, the plasma membrane from a slice of a serial section TEM dataset of first instar larva ventral nerve cord [16] of the *Drosophila melanogaster* was segmented (Fig. 1A, Supplementary Movie S1). The second example demonstrates the segmentation and separation of objects from 2D-LM images. Here, we segmented nuclei, their boundaries, and detected interfaces between adjacent nuclei from a high-throughput screen on cultured cells [17] (Fig. 1B, Supplementary Movie S2). The third and fourth examples demonstrate the segmentation of 3D EM and multicolor 3D LM datasets, where we segmented mitochondria from the mouse CA1 hippocampus [18] (Fig. 1C, Supplementary Movie S3) and inner hear cell cytoplasm, nuclei and ribbon synapses from mouse inner ear cochlea [19] (Fig. 1D, Supplementary Movie S4), respectively. In all cases, DeepMIB was able to achieve satisfactory results with modest time investment (see Supplementary Material for details). When necessary, the generated models can be further manually or semi-automatically proofread using MIB for quantitative analysis.

**Figure 1.**
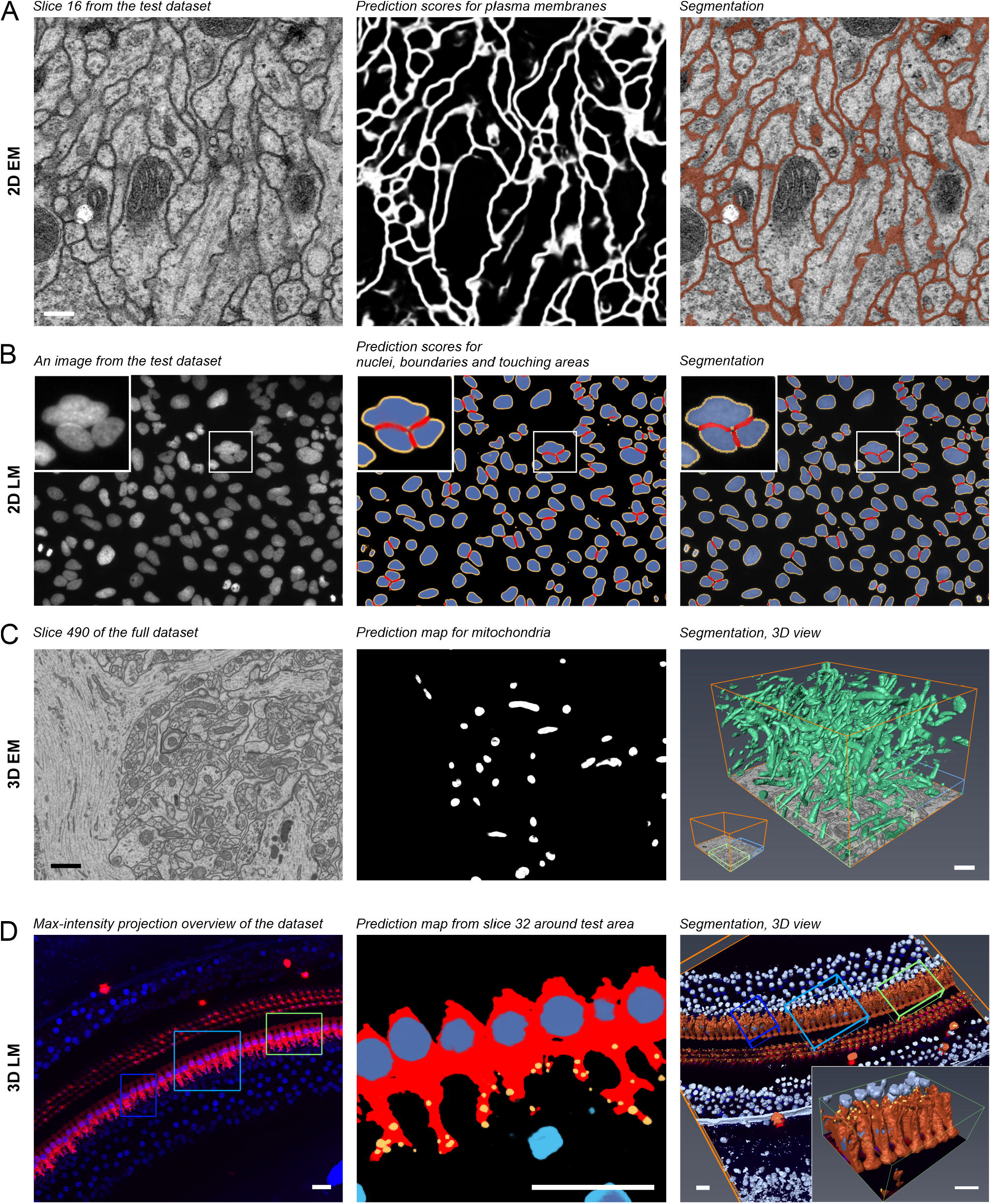
Test datasets and acquired segmentation results. (A) a middle slice of a serial section transmission electron microscopy dataset of the *Drosophila melanogaster* first instar larva ventral nerve cord supplemented with membrane prediction map and final segmentation of plasma membranes. (B) a random image of DNA channel from a high-throughput screen on human cultured osteocarcinoma U2OS cells (BBBC022 dataset, Broad BioImage Benchmark Collection) supplemented with prediction maps and final segmentation of nuclei, their boundaries (depicted in yellow), and interfaces between adjacent nuclei (depicted in red). The inset highlights a cluster of three nuclei. (C) a slice from a focus ion bean scanning electron microscopy dataset of the CA1 hippocampus region supplemented with prediction maps and 3D visualization of segmented mitochondria. In the right image, the blue box depicts area used for training and green box for testing and evaluation of the network performance. (D) a maximum intensity projection of 3D LM dataset supplemented with predictions and segmentation of inner hair cell located in the cochlea of the mouse inner ear. The inner hair cell cytoplasm depicted in vermillion, their nuclei in dark blue, and ribbon synapses in yellow. Nuclei of the surrounding cells are depicted in light blue. The light blue box indicates the area used for training, dark blue box for validation and green box (magnified in the inset) for testing and evaluation. The dataset was segmented using 3D U-net Anisotropic architecture, which was specially designed for anisotropic datasets. Scale bars, (A) 200 nm, (C) 1 μm, (D) 20 μm. (Scale bar for (B) not known). All presented examples are supplemented with movies (Supplementary Movie S1-4) and DeepMIB projects including datasets and trained network (Supplementary Material).

## Deep learning workflow

DeepMIB deep learning workflow comprises three main steps: preprocessing, training and prediction (Fig. 2a). Preprocessing requires images supplemented with the ground truth, which can be generated directly in MIB or using external tools [10, 11, 20, 21]. Typically, the application of deep learning to image segmentation requires large training sets. However, DeepMIB utilizes sets of 2D and 3D CNN architectures (U-Net [6, 8], SegNet [22]) that can provide good results already with only a few training data (starting from 2 to 10 images [13]), making it useful for small-scale projects too. To prevent overfitting of the network during training, the segmented images are randomly split into two sets, where the larger set (80-90%) is used for training and the smaller set for validation of the training process. DeepMIB is splitting the images automatically using configurable settings. When only one 3D dataset is available, it can be split into multiple subvolumes using the Chopping tool of MIB to dedicate one subvolume for validation. Alternatively, training can be done without validation. A modern GPU is essential for efficient training, but small datasets can still be trained using a central processing unit (CPU) only. Finally, in the prediction step, the trained networks are used to process unannotated datasets to generate prediction maps and segmentations.

**Figure 2.**
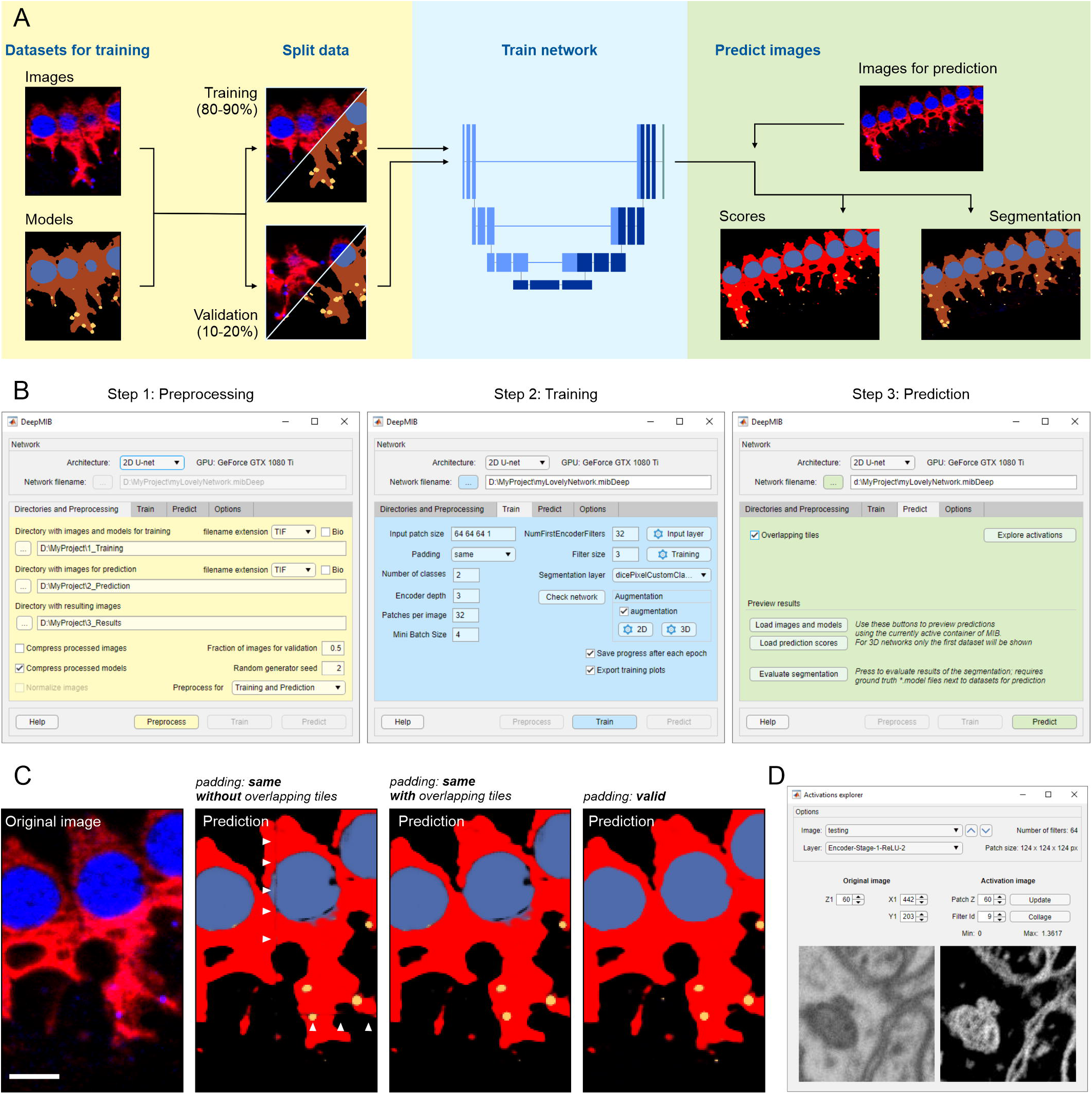
Schematic presentation of DeepMIB deep learning workflow. (A) a scheme of a typical deep learning workflow. (B) stepwise workflow of DeepMIB user interface for training or/and prediction. (C) comparison of different padding types and overlapping strategies during prediction. Description for used images is given in Fig. 1D. Arrowheads denote artefacts that may appear during prediction of datasets trained with the “same” padding parameter. (D) a snapshot of the GUI of the Activations explorer tool used for visualization of network features. Scale bar, 5 μm.

## User interface and training process

The user interface of DeepMIB (available from MIB *Menu->Tools->Deep learning segmentation*) consists of a single window with a top panel and several tabs arranged in a pipeline order and color-coded to help navigation (Fig. 2b). The process starts from choosing the network architecture from four options, 2D or 3D U-Net [6, 8], 3D U-Net designed for anisotropic datasets (Supplementary Fig. S1) or 2D SegNet [22], as preprocessing of datasets is determined according to the selected CNN. To improve usability and minimize data conversion, DeepMIB accepts both standard (*e.g.,* TIF, PNG) and microscopy (Bio-Formats [23]) image formats. The network training is done using the Train tab. DeepMIB automatically generates a network layout based on the most critical parameters that the user has to specify: the input patch size, whether to use convolutional padding, the number of classes and the depth of the network. With these parameters, the network layout can be tuned for specific dataset and available computational power. To improve end-results, data augmentation is a powerful method to expand the training set by using various transformations such as reflection, rotation, scaling, and shear. DeepMIB has separate configurable augmentation options for 2D and 3D networks. The training set can also be extended by filtering training images and their corresponding ground truth models using Elastic Distortion [24] filter of MIB (available from MIB *Menu->Image->Image filters*). For final fine-tuning, the normalization settings of the input layer and thorough tweaking of training parameters can be done. It is possible to store network checkpoints after each iteration and continue training from any of those steps.

## Prediction of new datasets

The prediction process is rather simple and requires only loading of a network file and preprocessing of images for prediction. Overlapping tiles option can be used to reduce edge artefacts during prediction (Fig. 2c). The results can be instantly checked in MIB and their quality can be evaluated against the ground truth with various metrics (*e.g.,* Jaccard similarity coefficient, F1 Score), providing that the ground truth model for the prediction dataset is available. In addition, Activations Explorer provides means to follow image perturbations inside the network and explore the network features (Fig. 2d).

## Summary

DeepMIB is included into MIB distribution, which is easy to install on any workstation or virtual machine preferably equipped with GPU. DeepMIB can train 2D and 3D networks for EM and LM for isotropic and anisotropic voxels. For a better understanding of the procedure and reducing the learning curve, DeepMIB has a detailed help section, online tutorials, and we provide workflows for all presented examples (Supplementary Material). At the moment, DeepMIB offers four CNNs, but as a future perspective to fulfill the needs of more experienced users, we are aiming to increase the list, provide a configuration tool for designing of own networks or import networks trained elsewhere.

## Acknowledgements

The research was supported by Biocenter Finland. We would like to thank Kuu Ikäheimo and Ulla Pirvola (the Auditory Physiology group, University of Helsinki) for kindly providing the inner ear dataset. The ISBI Challenge: Segmentation of neuronal structures in EM stacks [25] is acknowledged for evaluating the results of the 2D-EM segmentation. Dr. Konstantin Kogan (University of Helsinki) is thanked for helping with testing of MacOS version. Prof. Jussi Tohka and Mr. Ali Abdollahzadeh (University of Eastern Finland), and Dr. Helena Vihinen (Electron Microscopy Unit, Institute of Biotechnology, University of Helsinki) are thanked for the valuable comments on the manuscript.

## Author contributions

E.J. led the project, I.B. designed and programmed DeepMIB. I.B. and E.J wrote the manuscript.

## Supporting Information

**Supplementary Material. Details of installation details, online tutorials and example datasets.**

**Supplementary Figure S1. Schematic representation of the 3D U-Net Anisotropic architecture.** The architecture is based on a standard 3D U-net, where the 3D convolutions and the Max Pooling layer of the 1^st^ encoding level are replaced with the corresponding 2D operations (marked using “2D” label). To compensate, the similar swap is done for the last level of the decoding pathway. The scheme shows one of possible cases with patch size of 128×128×64×2 (height x width x depth x color channels), 2 depth levels, 32 first level filters and using “same” padding. This network architecture can be tweaked by modifying configurable parameters and it works best for anisotropic voxels with 1 x 1 x 2 (x, y, z) aspect ratio.

**Supplementary Movie S1. Final segmentation results of the test dataset of serial section transmission electron microscopy of the *Drosophila melanogaster* first instar larva ventral nerve cord**. The video is accompanying Fig. 1A.

**Supplementary Movie S2. Final segmentation of nuclei (blue), their boundaries (yellow) and interfaces between adjacent nuclei (red) for random images from a high-throughput screen on human cultured osteocarcinoma U2OS cells (BBBC022 dataset, Broad BioImage Benchmark Collection)**. The video is accompanying Fig. 1B.

**Supplementary Movie S3. Segmentation of mitochondria from the full focused ion beam scanning electron microscopy dataset of the CA1 hippocampus region.** The video is accompanying Fig. 1C.

**Supplementary Movie S4. A 3D LM dataset from mouse inner ear cochlea.** The shown dataset has two color channels: blue, CtBP2 staining of nuclei and ribbon synapses, and red, myosin 7a staining, highlighting inner and outer hair cells. The bottom slice, shown with the model represents the maximum intensity projection through the z-stack. The focus of the study was to segment the inner hair cells and their synapses thus the training and the validation sets were made around those cells omitting other cell type. The video is accompanying Fig. 1D.

