## Supplementary figures and images for "DeepMIB: User-friendly and open-source software for training of deep learning network for biological image segmentation"

### Supplementary Figure S1

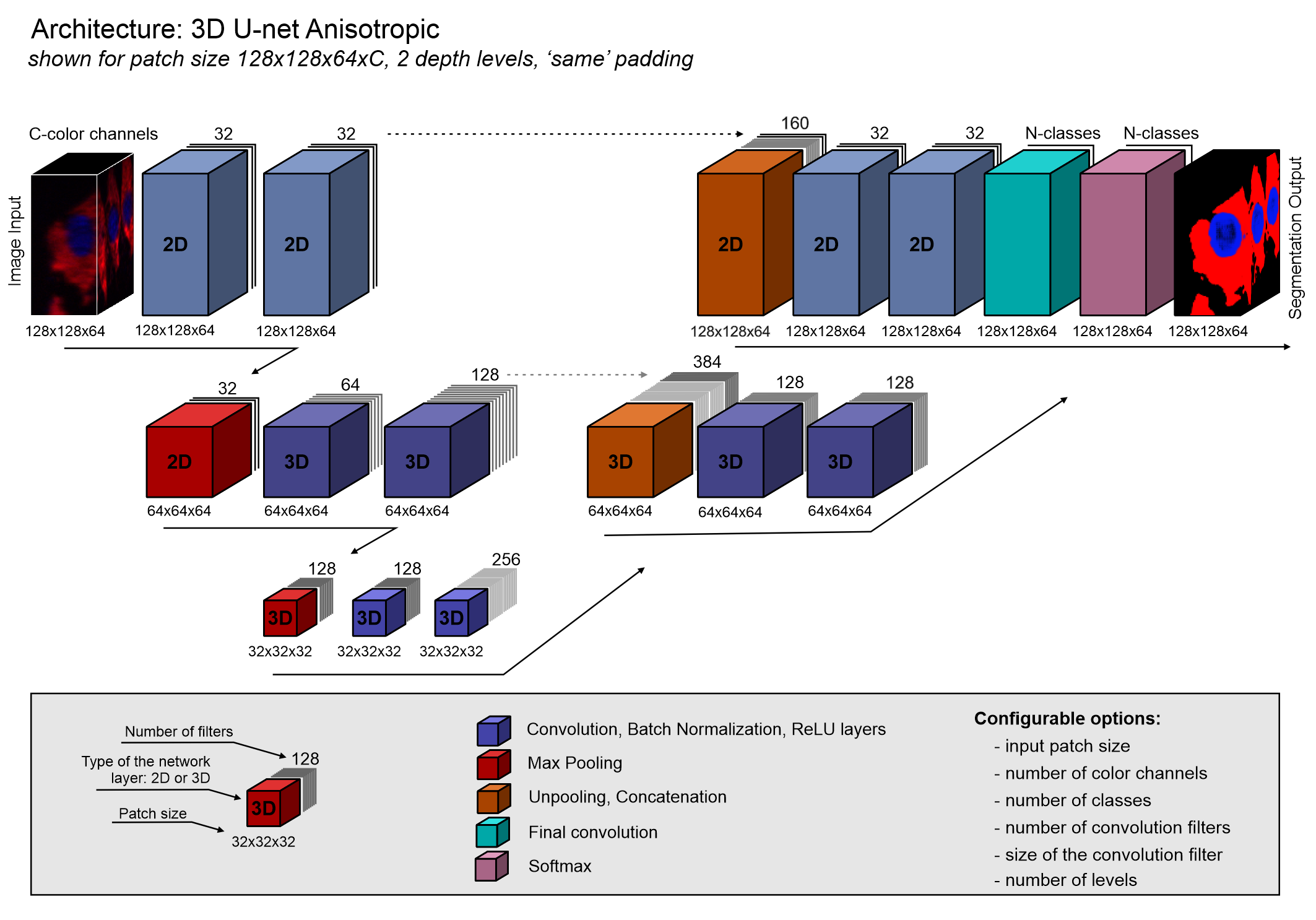
