## Supplementary Material for "DeepMIB: User-friendly and open-source software for training of deep learning network for biological image segmentation"

### Contents

Distribution versions and download links ... 1
System requirements ... 1
Installation details, system requirements and license ... 2
Demos and instructions to use ... 2
Workstation details ... 2
Supplementary information for Fig 1A ... 3
Supplementary information for Fig 1B ... 5
Supplementary information for Fig 1C ... 8
Supplementary information for Fig 1D ... 11
References: ... 14

#### Distribution versions and download links

DeepMIB is bundled with Microscopy Image Browser (MIB) [1] and installed automatically during installation of MIB (version 2.70 or newer).

MIB is provided in the following distribution packages:

- Matlab version, compatible with Windows, MacOS and Linux (*requires Matlab license*)
- Standalone, compiled for Windows (*does not require Matlab license*)
- Standalone, compiled for MacOS (*does not require Matlab license*)

Up-to-date code and new releases are available from:

- MIB website: http://mib.helsinki.fi/downloads.html
- GitHub: https://github.com/Ajaxels/MIB2 and https://github.com/Ajaxels/MIB2/releases

#### System requirements

**Matlab version of MIB+DeepMIB available for Windows, MacOS and Linux:**

- Matlab, version 9.7, R2019b (or newer)
- Computer Vision Toolbox, version 9.1, R2019b (or newer)
- Deep Learning Toolbox, version 13.0, R2019b (or newer)
- Image Processing Toolbox, version 11.0, R2019b (or newer)

Tested on Windows 10 (for Matlab R2019b and R2020a) and on MacOS (R2019b)

**Standalone MIB+DeepMIB, compiled for Microsoft Windows:**

- Windows, x64 bit

- All required libraries are automatically installed during the installation process

Tested on Windows 10, x64 (compiled with Matlab R2019b)

**Standalone MIB+DeepMIB, compiled for MacOS:**

- MacOS, x64 bit
- All required libraries are automatically installed during the installation process

Tested on MacOS, x64, Mojave, version 10.14.6 (compiled with Matlab R2019b)

#### Installation details, system requirements and license

Detailed installation guides are available on MIB website:

- Matlab version:
  - text description (http://mib.helsinki.fi/downloads_installation.html)
  - video demonstration (https://youtu.be/aGjys3nAmx0)
- Standalone version for Windows http://mib.helsinki.fi/downloads_installation_windows.html
- Standalone version for MacOS: http://mib.helsinki.fi/downloads_installation_macos.html

Installation time for Matlab version: 1-5 minutes depending on user experience with Matlab

Initial installation time for standalone version: ~7-15 minutes (depending on the network performance) for initial installation, which includes installation of required Matlab libraries, or ~30 seconds, when Matlab libraries are already installed on the system)

Microscopy Image Browser and DeepMIB are licensed under the GNU General Public License v2.

#### Demos and instructions to use

To start DeepMIB use: *MIB->Menu->Tools->Deep learning segmentation*

For better understanding of DeepMIB workflow, we released two online tutorials explaining GUI of DeepMIB and show all step required to achieve similar to presented in Fig. 1A and Fig. 1D results

- DeepMIB: How to train 2D U-Net for microscopy images (49 minutes), https://youtu.be/gk1GK_hWuGE
- DeepMIB: How to train 3D U-Net for microscopy images (34 minutes), https://youtu.be/U5nhbRODvqU

All details about the user interface are specified in DeepMIB Help system available upon press of the Help button in the right bottom corner of DeepMIB window

All presented results can be reproduced by loading the config file (mibCfg extension), which is accompanying each example from Fig 1 (*DeepMIB->Options tab->Config files->Load*). See below for detailed instructions and download links.

#### Workstation details

- Intel Xeon E5-2637 v2, @3.5GHz (2 cores), 192Gb RAM, NVidia GeForce GTX 1080 TI, 11Gb
- Windows 10, 64bit

- Matlab (Mathworks Inc.), R2019b with Deep Learning, Computer Vision, Image Processing Toolboxes
- Amira 2019.4 (Thermo-Fisher Scientific) was used for 3D rendering of models and supplementary movies 3 and 4.
- Microscopy Image Browser (MIB), version 2.70

#### Supplementary information for Fig 1A

Serial section Transmission Electron Microscopy dataset of the *Drosophila* first instar larva ventral nerve cord

**Dataset details**

The training and test datasets of the *Drosophila* first instar larva ventral nerve cord [2] obtained using serial section Transmission Electron Microscopy (ssTEM). The dataset was downloaded (http://brainiac2.mit.edu/isbi_challenge/home) from ISBI Challenge: Segmentation of neuronal structures in EM stacks [3].

The training set consisted of 30 8-bit annotated sections with dimensions (height x width) of 512x512 pixels and 1 color channel acquired with a resolution of 4x4x50 nm/voxel. The test set has the same dimensions but provided unannotated. Because the dataset has large anisotropic voxels we used 2D U-Net for its segmentation.

**Preparation of downloaded datasets for DeepMIB**

All image processing steps were done in MIB:

1. Load "train-volume.tif" to MIB. The contrast of the dataset is not normalized, i.e. some slices brighter that the others. We used the dataset as it is without contrast normalization (even though it is available in MIB) to test the robustness of the method

2. Load the model file: "train-labels.tif"; the model encodes cells with index 255 and membranes with index 0

3. Select material 255 (*cells*) in the Segmentation table, press *Alt+A* to highlight it as the Selection layer of MIB; invert the Selection layer (*Menu->Selection->Invert selection*) to highlight membranes.

4. Create a new model (*Menu->Models->New model*) and add a new material "*membranes*" (the "+" button in the Segmentation panel), press *Shift+A* to assign membranes to this new material of the model

5. Save dataset (*Menu->File->Save as*) to "MyProject\1_Training" directory using *AmiraMesh binary file sequence format*, which creates a sequence of 2D files; alternatively TIF format can be used.

6. Save model (*Menu->Models->Save model as*) to the same directory using **.model* format

7. Load "test-volume.tif" and save it to "MyProject\2_Prediction" directory using *AmiraMesh binary file sequence format*

**Network training**

**Network settings: 2D U-Net**

| | | | |
|---|---|---|---|
| Input patch size: | 508 508 1 1 | NumFirstEncoderFilters: | 64 |
| Padding: | valid | Filter size: | 3 |
| Number of classes: | 2 | Segmentation layer: | dicePixelCustomClassificaionLayer |

| | | | |
|---|---|---|---|
| Encoder depth: | 4 | Augmentation: | on |
| Patches per image: | 64 | Input layer normalization: | rescale-zero-one, Min=0, Max=255 |
| Mini batch size: | 1 | Validation fraction: | 0.15 |

**Augmentation settings**

| | | | |
|---|---|---|---|
| Fill value | 0 | RandScale: | 0.9 1.1 |
| RandXReflection: | on | RandXScale: | 1 1 |
| RandYReflection: | on | RandYScale: | 1 1 |
| RandRotation: | -15 15 | RandXShear: | -10 10 |
| | | RandYShear: | -10 10 |

**Training settings**

| | | | |
|---|---|---|---|
| SolverName | sgdm | LearnRateDropFactor: | 0.1 |
| MaxEpoch: | 50 | L2Regularization: | 0.0001 |
| Shuffle: | every-epoch | Momentum: | 0.9 |
| InitialLearnRate: | 0.005 | ValidationFrequency: | 400 |
| LearnRateSchedule: | piecewise | Plots: | training-progress |
| LearnRateDropPeriod: | 10 | | |

**Training progress plot:**

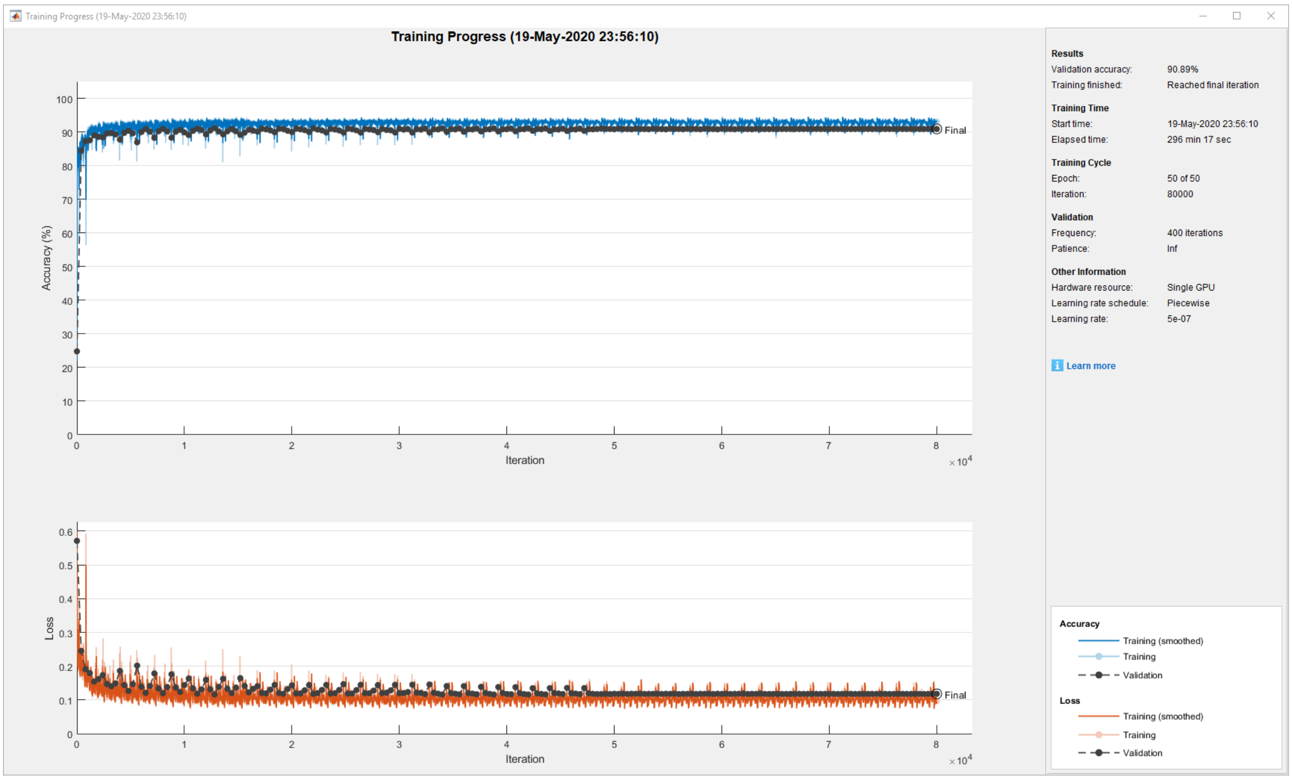


**Timing:** training: 4h 56min 17sec; prediction: 8.35 seconds for 30 images (0.28 image/sec)

**Evaluation scores:**

The presented scored were acquired from organizers of the ISBI Challenge: Segmentation of neuronal structures in EM stacks for the inverted model, where membranes are encoded with 0 (Exterior) and cells as 255.

Maximal foreground-restricted Rand score after thinning: 0.912748128
Maximal foreground-restricted information theoretic score after thinning: 0.969861473

**DeepMIB project and installation instructions:**

The full project directory is available via this link: http://mib.helsinki.fi/tutorials/deepmib/Figure1aFiles.zip

- Unzip the archive
- Start DeepMIB (MIB->Menu->Tools->Deep Learning segmentation)
- Load the config "*valid_508px_64pat.mibCfg*" file (DeepMIB->Options tab->Config files->Load)

#### Supplementary information for Fig 1B

DNA channel of a high-throughput chemical screen on U2OS cells

**Dataset details**

The dataset (BBBC039) was acquired from Broad Bioimage Benchmark Collection [4] (https://data.broadinstitute.org/bbbc/BBBC039). It consisted of 197 16-bit annotated images (3 empty images without cells were removed) with dimensions (height x width) of 520 x 696 pixels and 1 color channel. The dataset was randomly split into the training (143 images) and prediction sets (54 images). Training was done using 2D U-Net.

The representative slice, shown in Fig. 1b with original filename: IXMtest_J08_s2_w1C146DB1C-05B3-49EF-9C62-1185FD9897AC.tif was selected as the representative one due to multiple touching nuclei.

**Preparation of downloaded datasets for DeepMIB**

All image processing steps were done in MIB:

1. Load files from masks.zip to MIB

2. Delete channel 2 and channel 3 (*Menu->Image->Color channels->Delete channel*)

3. Save dataset in TIF format as model.tif

4. Load image files from images.tif

5. Load model.tif file as a model (*Menu->Models->Load model*)

6. Use the Image Arithmetics tool (*Menu->Image->Tools for images->Image arithmetics*) to generate model with 3 objects: nuclei, boundaries, and touching boundaries:

Set Input and Output variables: **O**

Run the following expression:

```
% define strel element
se = zeros([5 5],'uint8');
se(3,3) = 1;
se = bwdist(se);
se = uint8(se <= max(2));
% generate masks for each material
M1 = zeros(size(O), 'uint8');
M2 = zeros(size(O), 'uint8');
M3 = zeros(size(O), 'uint8');
M1(O==1) = 1;
```

```
M2(O==2) = 1;
M3(O==3) = 1;
% assign nuclei
O(O>0) = 1;
% assign boundary
O(M1-imerode(M1, se) == 1) = 2;
O(M2-imerode(M2, se) == 1) = 2;
O(M3-imerode(M3, se) == 1) = 2;
% assign touching boundaries
M = imdilate(M1, se);
O(M2 & M) = 3;
O(M3 & M) = 3;
M = imdilate(M2, se);
O(M1 & M) = 3;
O(M3 & M) = 3;
M = imdilate(M3, se);
O(M1 & M) = 3;
O(M2 & M) = 3;
```

7. Rename materials to: Nuclei, Boundary, TouchingBoundary

8. Delete empty slices (*Menu->Dataset->Slice->Delete slice*): **69** (IXMtest_F13_s7_w13C1B1D8C-293E-454F-B0FD-6C2C3F9F5173.tif), **139** (IXMtest_L01_s2_w1E5038251-DBA3-44D0-BC37-E43E2FC8C174.tif), **146** (IXMtest_L10_s6_w12D12D64C-2639-4CA8-9BB4-99F92C9B7068.tif)

9. Save dataset (*Menu->File->Save as*) in TIF format to a new directory using: "*use original filenames*" and "*2D sequence*" as options.

10. Save model to the same directory using *.model format (*Menu->Models->Save model as*)

11. To randomly split dataset for training and prediction we used Rename and shuffle tool of MIB (*Menu->File->Rename and shuffle->Rename and shuffle*): random seed: 0, number of output directories: 4, include model: on. We generated 4 sets: 3 of those sets (143 images in total) were combined into *1_Training* directory and 4$^{th}$ set (54 images) to *2_Prediction*.

**Network training**

**Network settings: 2D U-Net**

| | | | |
|---|---|---|---|
| Input patch size: | 252 252 1 1 | NumFirstEncoderFilters: | 32 |
| Padding: | valid | Filter size: | 3 |
| Number of classes: | 4 | Segmentation layer: | dicePixelCustomClassificaionLayer |
| Encoder depth: | 3 | Augmentation: | on |
| Patches per image: | 32 | Input layer normalization: | zscore |
| Mini batch size: | 4 | Validation fraction: | 0.25 |

**Augmentation settings**

| | | | |
|---|---|---|---|
| Fill value | 0 | RandScale: | 0.9 1.1 |
| RandXReflection: | on | RandXScale: | 1 1 |
| RandYReflection: | on | RandYScale: | 1 1 |
| RandRotation: | -20 20 | RandXShear: | -10 10 |
| | | RandYShear: | -10 10 |

**Training settings**

| | | | |
|---|---|---|---|
| SolverName | sgdm | LearnRateDropFactor: | 0.1 |
| MaxEpoch: | 50 | L2Regularization: | 0.0001 |

| | | | |
|---|---|---|---|
| Shuffle: | every-epoch | Momentum: | 0.9 |
| InitialLearnRate: | 0.005 | ValidationFrequency: | 400 |
| LearnRateSchedule: | piecewise | Plots: | training-progress |
| LearnRateDropPeriod: | 10 | | |

**Training progress plot:**

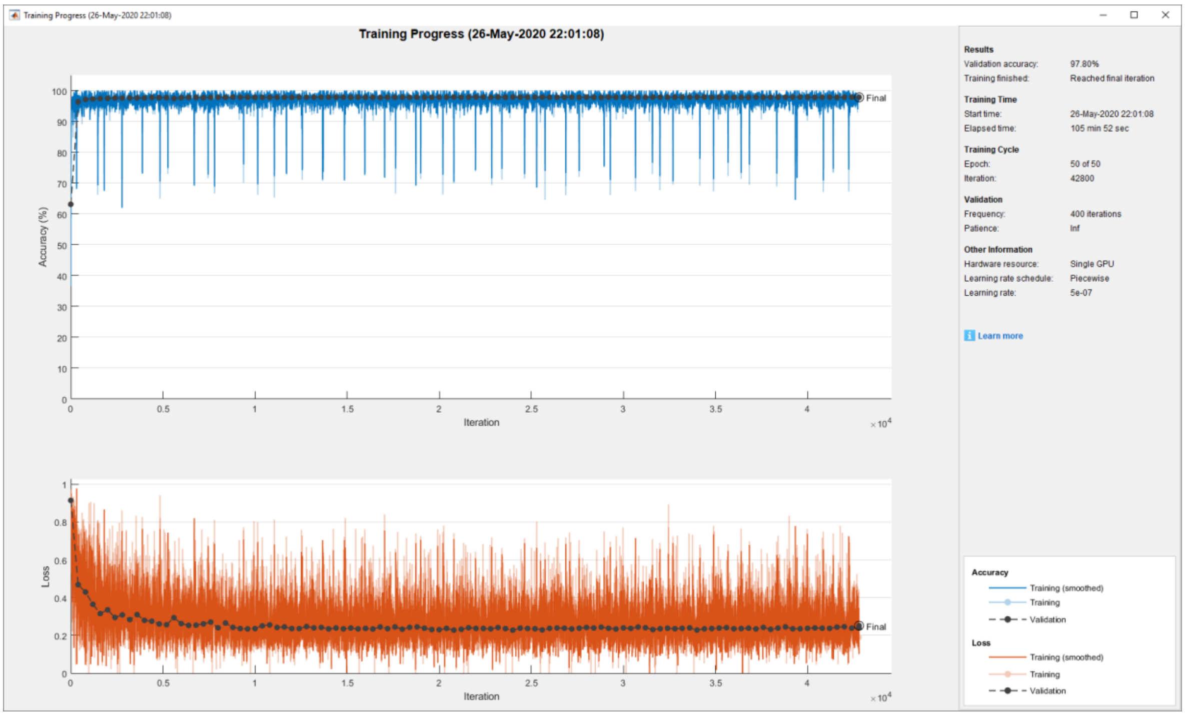


**Timing:** training: 1h 45min 52sec; prediction: 38.0 seconds for 54 images (0.70 image/sec)

**Evaluation scores:**

Dataset metrics

| GlobalAccuracy | MeanAccuracy | MeanIoU | WeightedIoU | MeanBFScore |
|---|---|---|---|---|
| 0.977084254 | 0.860487336 | 0.73808 | 0.960118779 | 0.935740352 |

Class metrics

| | Accuracy | IoU | MeanBFScore |
|---|---|---|---|
| *Exterior* | 0.992293498 | 0.986306968 | 0.998579 |
| *Nuclei* | 0.945271161 | 0.921782612 | 0.99788 |
| *Boundary* | 0.819574456 | 0.63776319 | 0.996986 |
| *TouchingBoundary* | 0.684810227 | 0.406470921 | 0.749517 |

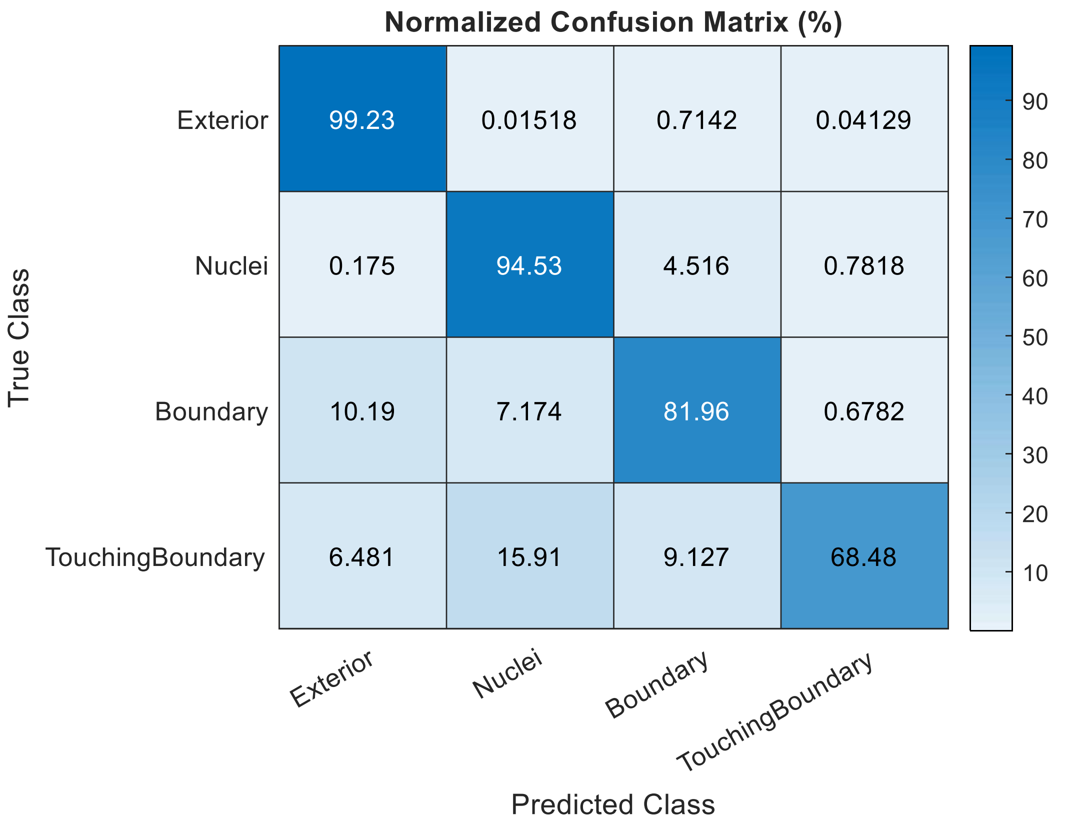


**DeepMIB project and installation instructions:**

The full project directory is available via this link: http://mib.helsinki.fi/tutorials/deepmib/Figure1bFiles.zip

- Unzip the archive
- Start DeepMIB (MIB->Menu->Tools->Deep Learning segmentation)
- Load the config “*valid_252px_32patches_50ep.mibCfg*” file (DeepMIB->Options tab->Config files->Load)

#### Supplementary information for Fig 1C

FIB-SEM electron microscopy dataset of the CA1 hippocampus.

**Dataset details**

The dataset was collected by Graham Knott and Marco Cantoni. The full dataset as well as two sub-volumes with the segmented ground truth are publicly available from the CVLAB, EPFL website: https://www.epfl.ch/labs/cvlab/data/data-em

The training set [5] consisted of 165 8-bit annotated sections with dimensions (height x width) of 768x1024 pixels and 1 color channel acquired with isotropic resolution of 5x5x5 nm/voxel (blue box in Fig. 1c). The test dataset (green box in Fig. 1c), which was used for evaluation of the network performance, has the same dimensions. The image in Fig. 1c shows prediction of mitochondria for the full 3D-EM dataset with dimensions: 2048 x 1536 x 1065 pixels. The training was done using 3D U-Net.

The representative slice, shown in Fig. 1c comes from the middle of the large dataset and has index 490.

**Preparation of downloaded datasets for DeepMIB**

All image processing steps were done in MIB:

1. Load training sub-volume (*training.tif*) to MIB

2. Load ground truth for training sub-volume (*training_groundtruth.tif*) as model.

3. Because mitochondria are encoded with index 255 we did the following operation:

- selected material 255, in the Segmentation table of MIB
- copied it to the Selection layer (*Alt+A* key shortcut)
- created a new model (*Menu->Models->New model*)
- added a new material "*mitochondria*" (the "+" button in the Segmentation panel)
- moved the Selection layer to this new material (*Shift+A* key shortcut)

4. Save the model into the same directory in *.model format

5. Repeat steps: 1-4 for the prediction dataset, but save it to *2_Prediction* directory

**Network training**

**Network settings: 3D U-Net**

| | | | |
|---|---|---|---|
| Input patch size: | 128 128 128 1 | NumFirstEncoderFilters: | 32 |
| Padding: | valid | Filter size: | 3 |
| Number of classes: | 2 | Segmentation layer: | dicePixelCustomClassificaionLayer |
| Encoder depth: | 2 | Augmentation: | on |
| Patches per image: | 256 | Input layer normalization: | zerocenter |
| Mini batch size: | 1 | Validation fraction: | 0 (validation was not used) |

**Augmentation settings**

| | | | |
|---|---|---|---|
| RandXReflection: | on | Fraction: | 0.8 |
| RandYReflection: | on | Rotation90: | on |
| RandZReflection: | on | ReflectedRotation90: | on |

**Training settings**

| | | | |
|---|---|---|---|
| SolverName | sgdm | LearnRateDropFactor: | 0.1 |
| MaxEpoch: | stopped at 176 | L2Regularization: | 0.0001 |
| Shuffle: | every-epoch | Momentum: | 0.9 |
| InitialLearnRate: | 0.005 | ValidationFrequency: | 400 |
| LearnRateSchedule: | none | Plots: | training-progress |
| LearnRateDropPeriod: | 10 | | |

##### Training progress plot:

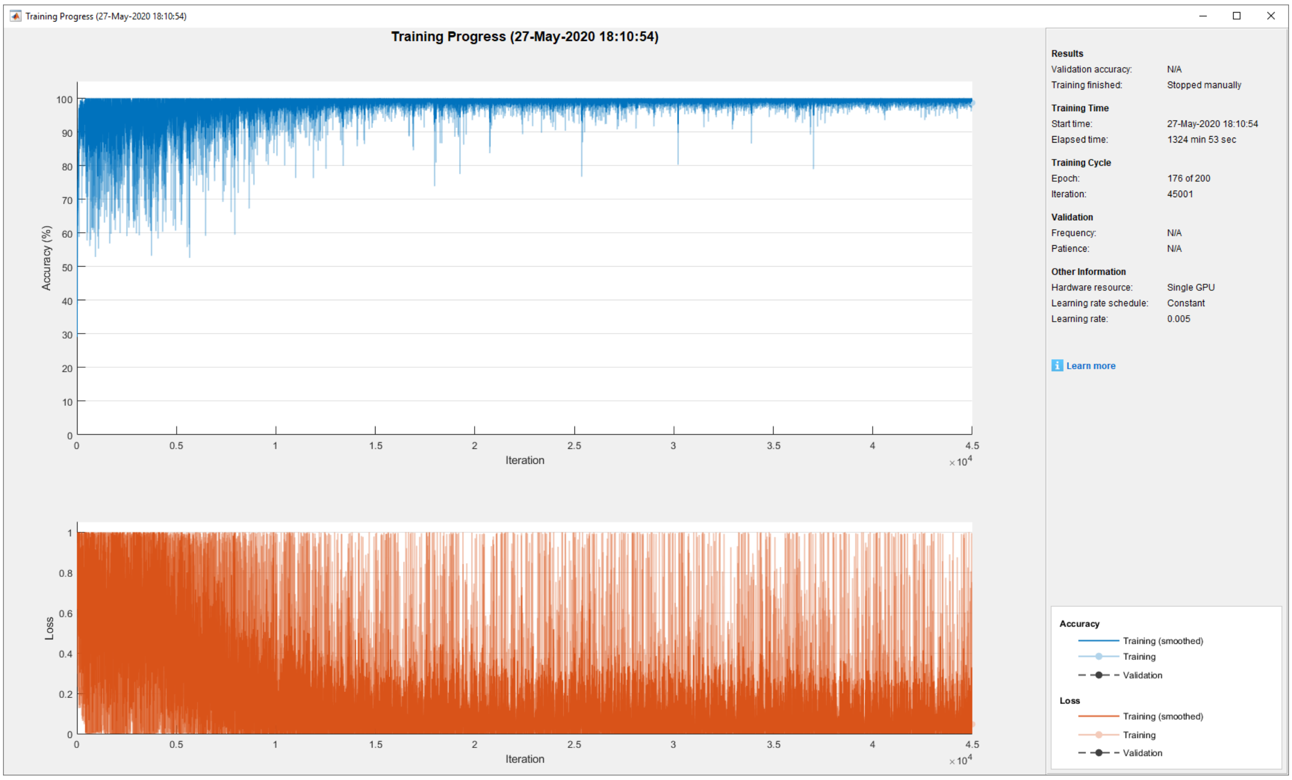


##### Timing:

- training: 22 hours 4 mins 53 sec
- prediction: 2 mins 12 seconds for the test volume; 58 mins for the full dataset

##### Evaluation scores:

Dataset metrics

| GlobalAccuracy | MeanAccuracy | MeanIoU | WeightedIoU | MeanBFScore |
|---|---|---|---|---|
| 0.990589859 | 0.955705329 | 0.91403 | 0.982031557 | 0.958768537 |

Class metrics

| | Accuracy | IoU | MeanBFScore |
|---|---|---|---|
| *Exterior* | 0.994733515 | 0.990108517 | 0.983299 |
| *Mitochondria* | 0.916677143 | 0.837958305 | 0.934238 |

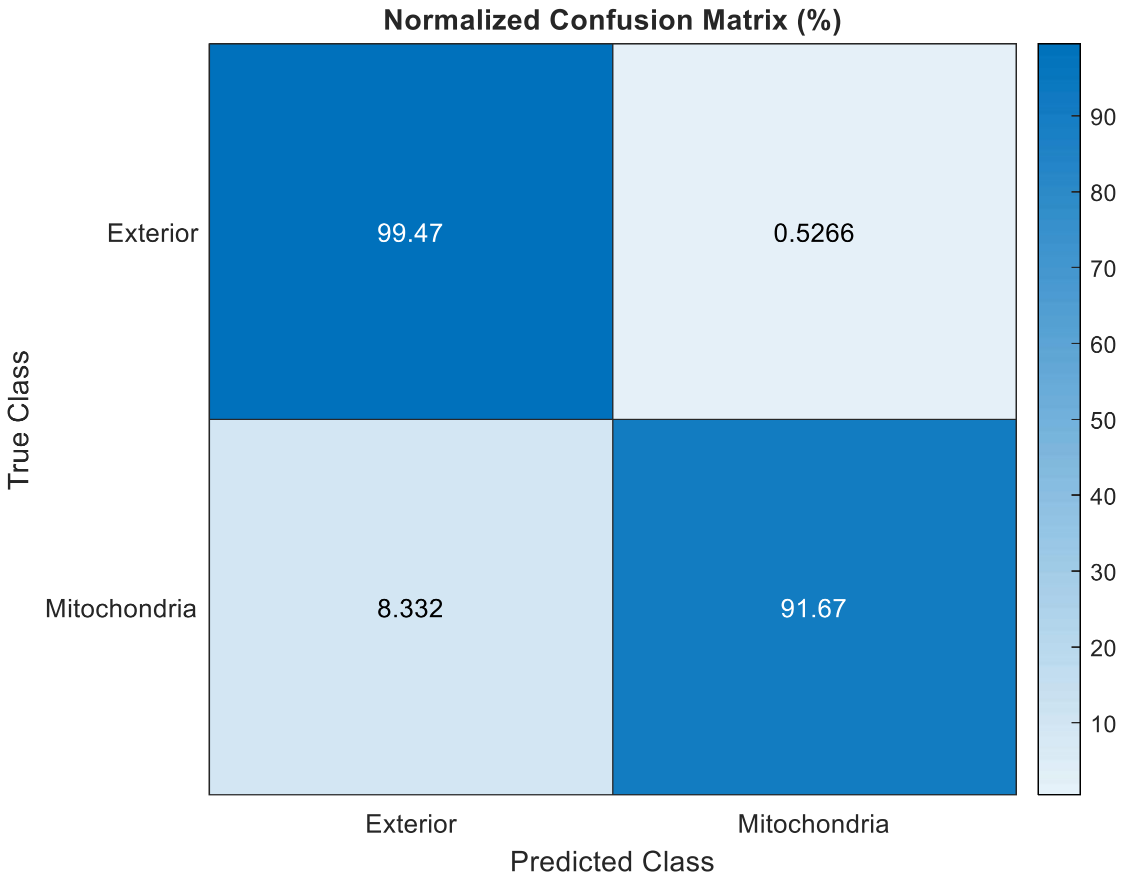


**DeepMIB project and installation instructions:**

The full project directory is available via this link: http://mib.helsinki.fi/tutorials/deepmib/Figure1cFiles.zip

- Unzip the archive
- Start DeepMIB (MIB->Menu->Tools->Deep Learning segmentation)
- Load the config "*NoValidation_valid_Aug_128px_256pat.mibCfg*" file (DeepMIB->Options tab->Config files->Load)

The archive includes the trained network and only two small subvolumes. The full dataset is available directly from the CVLAB website.

#### Supplementary information for Fig 1D

Inner hair cells of mouse inner ear

**Dataset details**

The dataset was kindly provided by Kuu Ikäheimo and Ulla Pirvola from the Auditory Physiology group, University of Helsinki. The sample preparation and imaging procedure are described in [6].

The dataset has 70 16-bit sections acquired with voxel size of 0.1625 x 0.1625 x 0.3 µm and dimensions (height x width) 2048 x 2048 and 2 color channels:

- channel 1: (blue), CtBP2 staining of nuclei and ribbon synapses
- channel 2: (red), myosin 7a staining, highlighting inner and outer hair cells

The dataset was processed as described below into four datasets: for training, validation, testing and full dataset for prediction:

- training dataset (height x width x depth x colors): 392 x 438 x 70 x 2
- validation dataset (height x width x depth x colors): 279 x 234 x 70 x 2
- test dataset (height x width x depth x colors): 273 x 367 x 70 x 2
- full dataset (height x width x depth x colors): 2048 x 2048 x 70 x 2

The training was done using 3D U-Net Anisotropic (Supplementary Fig. 1), which is optimal for volumes with anisotropic voxels

**Image processing**

1. Load the provided dataset in MIB

2. Save dataset using Amira Mesh format (*full dataset*)

3. Crop the dataset in three various areas and use those crops as volumes for training (*40x_OrganCorti_Training _2col.am*), validation (*40x_OrganCorti_Validation _2col.am*) and testing (*40x_OrganCorti_Testing _2col.am*) and save them

4. Segment the training, validation and testing volumes in MIB to create ground truth models

**Network training**

**Network settings: 3D U-Net Anisotropic**

| | | | |
|---|---|---|---|
| Input patch size: | 136 136 64 2 | NumFirstEncoderFilters: | 32 |
| Padding: | same | Filter size: | 3 |
| Number of classes: | 5 | Segmentation layer: | dicePixelCustomClassificaionLayer |
| Encoder depth: | 3 | Augmentation: | on |
| Patches per image: | 96 | Input layer normalization: | zerocenter |
| Mini batch size: | 1 | Validation fraction: | 0.5 |

**Augmentation settings**

| | | | |
|---|---|---|---|
| RandXReflection: | on | Fraction: | 0.8 |
| RandYReflection: | on | Rotation90: | on |
| RandZReflection: | on | ReflectedRotation90: | on |

**Training settings**

| | | | |
|---|---|---|---|
| SolverName | sgdm | LearnRateDropFactor: | 0.1 |
| MaxEpoch: | 120 | L2Regularization: | 0.0001 |
| Shuffle: | every-epoch | Momentum: | 0.9 |
| InitialLearnRate: | 0.005 | ValidationFrequency: | 400 |
| LearnRateSchedule: | none | Plots: | training-progress |
| LearnRateDropPeriod: | 10 | | |

**Training progress plot:**

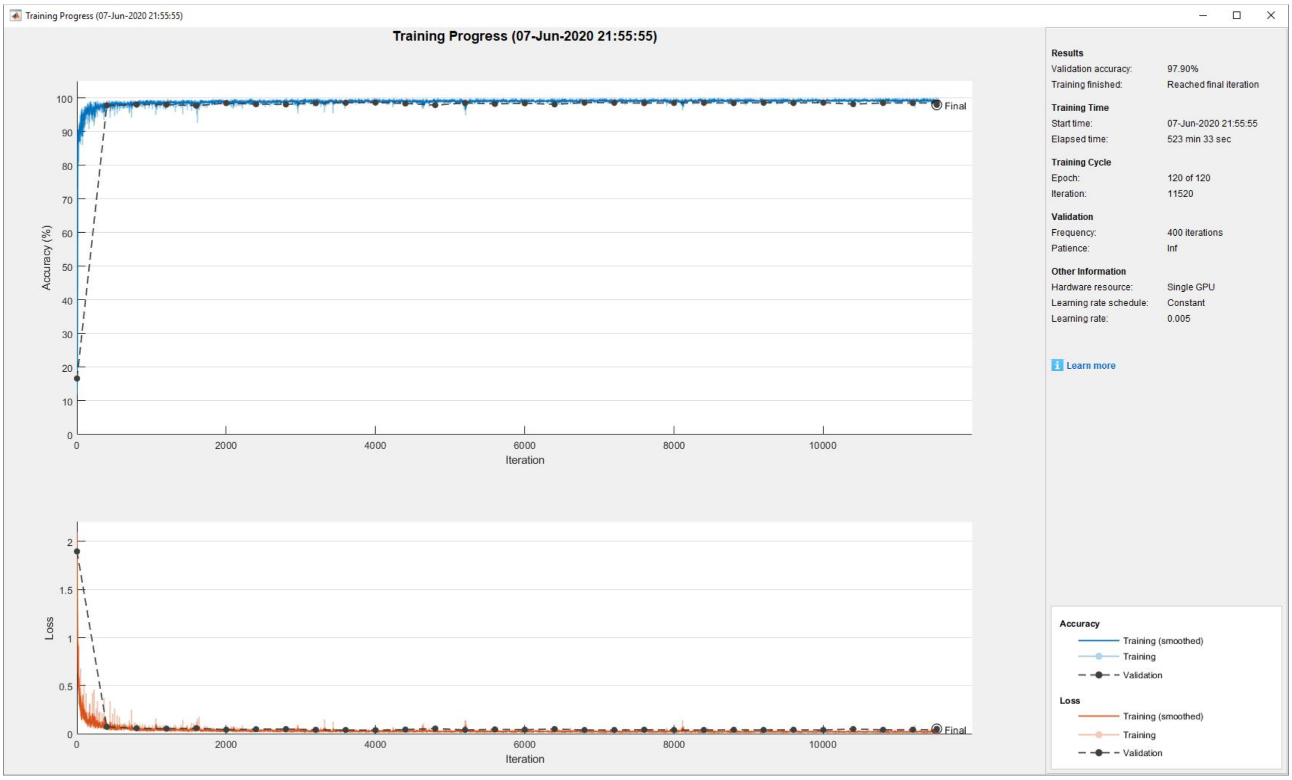


**Timing:**

- training: 8 hours 43 mins 33 sec
- prediction (with overlapping tiles): 20 seconds for the test volume; 9 mins 19 sec for the full dataset

**Evaluation scores:**

Dataset metrics

| GlobalAccuracy | MeanAccuracy | MeanIoU | WeightedIoU | MeanBFScore |
|---|---|---|---|---|
| 0.981958887 | 0.9424709 | 0.87389 | 0.965212776 | 0.976557523 |

Class metrics

| | Accuracy | IoU | MeanBFScore |
|---|---|---|---|
| *Exterior* | 0.991696108 | 0.982727078 | 0.996353 |
| *Nuclei* | 0.990488582 | 0.882292562 | 0.926482 |
| *Cells* | 0.965181014 | 0.933074763 | 0.993509 |
| *Syn* | 0.944073047 | 0.76847526 | 0.994073 |
| *NucleiOthers* | 0.82091575 | 0.802885266 | 0.97237 |

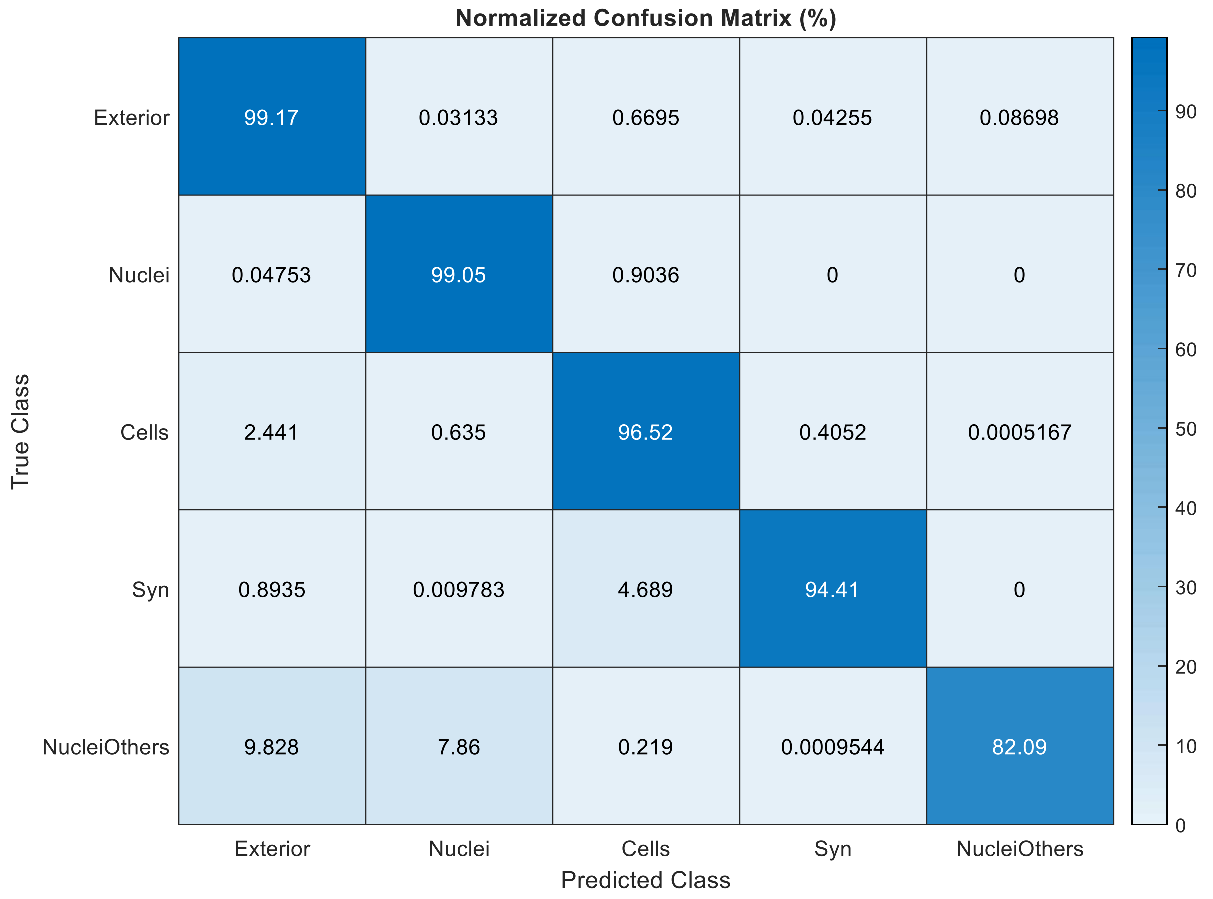


**DeepMIB project and installation instructions:**

The project directory (without the full dataset) is available via this link:
http://mib.helsinki.fi/tutorials/deepmib/Figure1dFiles.zip

- Unzip the archive
- Start DeepMIB (MIB->Menu->Tools->Deep Learning segmentation)
- Load the config "*InnerEar3D_Hybrid_Same_136x64px_120ep.mibCfg*" file (DeepMIB->Options tab->Config files->Load)
